# Drug Sensitivity Prediction From Cell Line-Based Pharmacogenomics Data: Guidelines for Developing Machine Learning Models

**DOI:** 10.1101/2021.04.09.439076

**Authors:** Hossein Sharifi-Noghabi, Soheil Jahangiri-Tazehkand, Petr Smirnov, Casey Hon, Anthony Mammoliti, Sisira Kadambat Nair, Arvind Singh Mer, Martin Ester, Benjamin Haibe-Kains

## Abstract

The goal of precision oncology is to tailor treatment for patients individually using the genomic profile of their tumors. Pharmacogenomics datasets such as cancer cell lines are among the most valuable resources for drug sensitivity prediction, a crucial task of precision oncology. Machine learning methods have been employed to predict drug sensitivity based on the multiple omics data available for large panels of cancer cell lines. However, there are no comprehensive guidelines on how to properly train and validate such machine learning models for drug sensitivity prediction. In this paper, we introduce a set of guidelines for different aspects of training gene expression-based predictors using cell line datasets. These guidelines provide extensive analysis of the generalization of drug sensitivity predictors, and challenge many current practices in the community including the choice of training dataset and measure of drug sensitivity. Application of these guidelines in future studies will enable the development of more robust preclinical biomarkers.

## INTRODUCTION

Cancer is a complex genetic disease. Due to the heterogeneous nature of tumors, the treatment of cancer is very challenging. Precision oncology aims to tailor the therapies according to the genomic profile of the tumor. Pharmacogenomics, a crucial component of precision oncology, promises to utilize the genomic landscape of each individual patient to find the most effective treatment options [5–7]. However, it still has limited clinical utility [8] and the availability of clinical pharmacogenomics datasets is limited by a lack of public access and small size, both in terms of patient cohorts and investigated therapies for the few publicly available datasets. As a result, resources such as cancer cell lines [1–4,9–11], patient-derived xenografts (PDX) [12,13], or organoids [14] are being employed in pharmacogenomics to decipher drug sensitivity prediction. Although these preclinical resources do not fully recapitulate the inter- and intra-tumor heterogeneity of cancer, they act as proxies for patient tumors and provide larger dataset, usually screened with hundreds or thousands of drugs separately or in combination along with multi-omics characterization [15].

Due to the complexity of generating pharmacogenomics datasets, discrepancies can even exist across cell line datasets [11,16–22]. However, recent efforts such as the PharmacoDB project (pharmacodb.ca) [15], the ORCESTRA platform (orcestra.ca) [23], and CellMinerCDB [24] aimed at standardizing, and integrating different preclinical pharmacogenomics datasets to improve downstream machine learning modeling. The data-rich nature of preclinical pharmacogenomics datasets has paved the way for the development of machine learning approaches to predict drug sensitivity *in vitro* and *in vivo* [25–27]. These computational approaches range from simple linear regression models [28,29] Lasso [30], and Elastic Net [31] to Random Forest [32], kernel-based models [33–36], highly non-linear models based on Deep Neural Networks [37–44], and most recently, reinforcement learning [45], few-shot learning [46], and multi-task learning [47]. These methods often take gene expression as input and predict the area above/under the dose-response curve (AAC/AUC) or half-maximal inhibitory concentration (IC50), the concentration of the drug that reduces the viability by 50%.

While machine learning for pharmacogenomics is a promising direction [25], existing guidelines are based on a single pharmacogenomics dataset [48] or based on benchmarking different methods without considering technical differences between molecular profiles or drug screening assays across different datasets [26]. We believe that there is a need for comprehensive guidelines based on multiple uniformly processed datasets on how to properly train and evaluate drug sensitivity predictors. In this study, we conduct a systematic and comprehensive analysis based on RNA-seq data as the input (gene expression-based models) and different measures of drug sensitivity such as AAC and IC50 as the output. We employ univariable modeling (using prospective biomarkers) and multivariable modeling (using state-of-the-art machine learning methods) to investigate generalization in drug sensitivity prediction. We consider two common machine learning paradigms: within-domain analysis and cross-domain analysis. In within-domain analysis, models are trained and tested on the same dataset via cross-validation which means train and test data are from the same distribution. In cross-domain analysis, models are trained and tested on different cell line datasets to investigate generalization capability. We also examine the effect of an analysis choice first proposed by [1], to separate the data for cell lines originating from hematopoietic cancers and solid tumours on the ability to learn predictors of drug sensitivity.

As a result of this study, we provide guidelines, which we refer to as PGx Guidelines (Figure 1), on the following questions:

**Figure 1.**
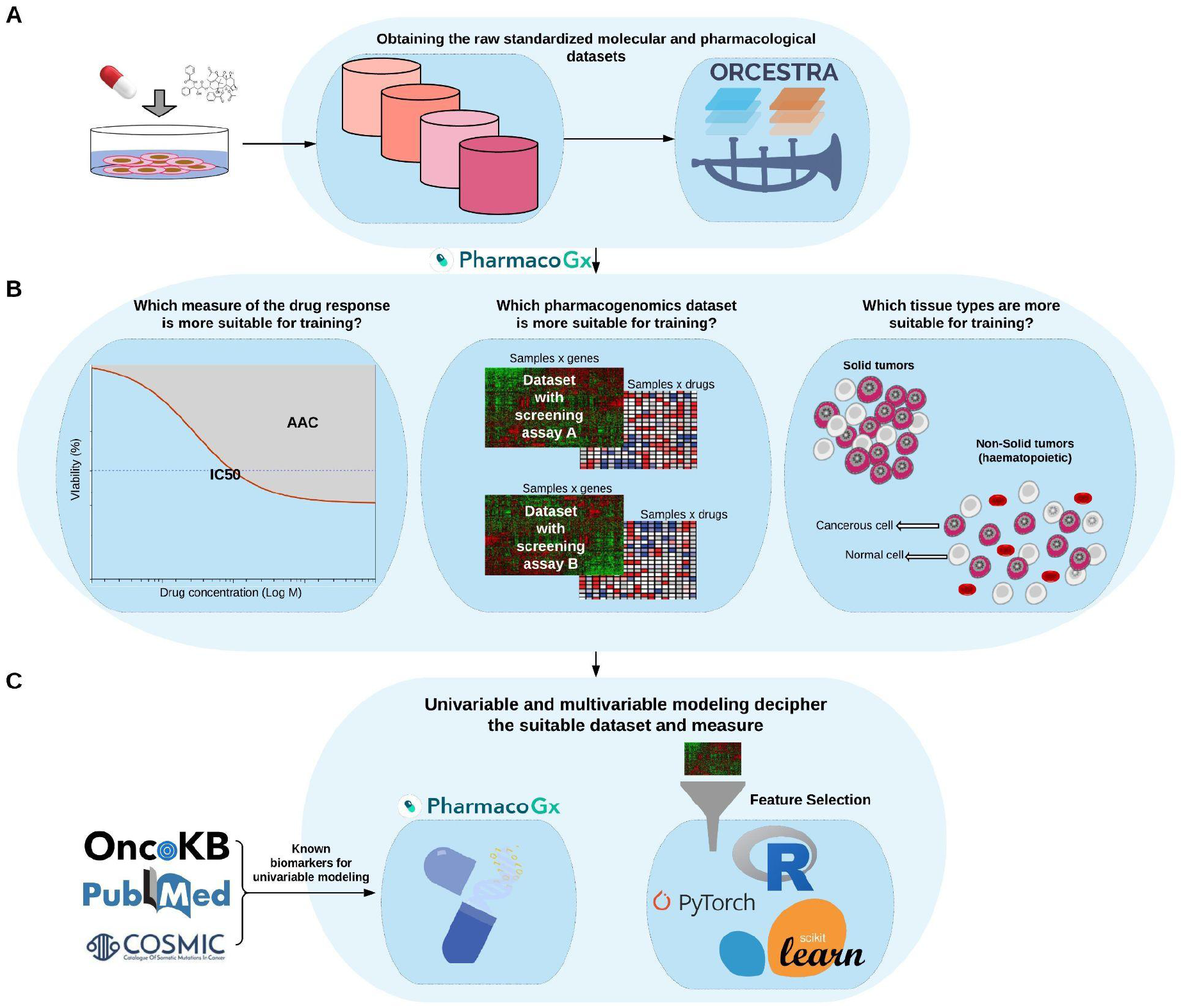
Schematic overview of the PGx Guidelines; A) The raw pharmacogenomics datasets are obtained from the ORCESTRA platform. B) The molecular (RNA-seq) and pharmacological profiles (Area Above dose-response Curve--AAC) are obtained for each dataset via the PharmacoGx package. Finally, C) univariable modeling based on prospective biomarkers from the literature is performed via the PharmacoGx package, and multivariable modeling using protein coding genes and feature selection is performed via different packages.

1. Which dataset(s) and measure(s) of drug sensitivity are best for training predictors?
2. How much does the performance of pharmacogenomics methods change when moving from within-domain analysis to cross-domain analysis?
3. What is the impact of non-solid tumors on the performance of drug sensitivity predictors?

We argue that it is necessary to evaluate generalization of cell line-based predictors first on cell line datasets before employing them on PDX or patient data, and therefore we focus on cell line datasets in this study. We believe that the PGx Guidelines will lead to the development of more accurate and more generalizable machine learning models for drug sensitivity prediction from pharmacogenomics data and will contribute toward the goal of extending the benefits of precision oncology to a wider range of patients.

## METHODS

### Drug Sensitivity Metrics

The datasets analyzed in this study combine molecular profiling of cancer cell lines with high throughput screening for drug sensitivity. For each drug-cell pair investigated in a dataset, cell viability at several increasing doses of the drug was measured and compared to an untreated control, to obtain % viability values. To learn predictors of drug response, it is desirable to obtain a single number summarizing a particular cell line’s sensitivity to a drug treatment (which can then be used as a label in fitting predictive models from the molecular features). We study two different summary measures: the Area Above the Curve (AAC) and the half maximal inhibitory concentration (IC50). Both of these measures are derived by first fitting a Hill Curve model to the dose-response data. To ensure consistency in the inference method, we fit a 3 parameter Hill Curve to all the datasets, using the *logLogisticRegression* function in the PharmacoGx R package, as described previously [17,49]. The AAC is then the area above the curve, integrated from the lowest to highest measured concentration, normalized to the concentration range. The IC50 is the concentration at which the curve crosses 50% viability. Some curves estimated in the data never cross this 50% threshold, and therefore the IC50 does not exist for many experiments where the AAC can be calculated. In this paper, both these values were calculated using methods implemented in the PharmacoGx package [49].

### Datasets

We employed the following pan-cancer datasets (Table 1):

**Table 1.** Characteristics of the studied datasets

| Characteristic | CTRPv2 | GDSCv2 | GDSCv1 | gCSI |
| --- | --- | --- | --- | --- |
| # Drugs | 544 | 190 | 343 | 16 |
| # Cell Lines | 821 | 328 | 427 | 334 |
| # Tissue Types | 25 | 27 | 28 | 22 |
| # Genes (# Protein Coding) | 60662 (19957) | 60662 (19957) | 60662 (19957) | 60662 (19957) |
| Gene Expression Assay | RNA-seq ** | RNA-seq | RNA-seq | RNA-seq |
| Sensitivity Assay | CellTiter Glo | CellTiter Glo | Syto60 | CellTiter Glo |
| Usage | Training | Test | Training | Test |

\* Obtained from the ORCESTR platform. Numbers are dependent to the preprocessing method for the RNA-seq
\*\* Gene expression for CTRPv2 were extracted from the CCLE dataset.

- The Cancer Therapeutics Response Portal (CTRPv2) [1,2]
- The Genentech Cell Line Screening Initiative (gCSI) [10,11]
- The Genomics of Drug Sensitivity in Cancer (GDSCv1 and GDSCv2) [3,4]

We obtained these datasets in the format of PharmacoSet (PSet) which is an R-based data structure that aids in reproducible research for drug sensitivity prediction. PSets are obtained via the ORCESTRA platform (orcestra.ca) [23]. The molecular profiles (RNA-seq) were preprocessed via Kallisto 0.46.1 [50] using GENCODE v33 transcriptome as the reference and the pharmacological profiles (AAC and IC50) were preprocessed and recomputed via PharmacoGx package [49]. In this paper, we focused on 11 drugs in common between these datasets including: Bortezomib, Entinostat, Sirolimus, Docetaxel, Gemcitabine, Crizotinib, Lapatinib, Vorinostat, Erlotinib, Paclitaxel, and Pictilisib. These datasets have missing values for different samples and given a specific drug, the number of available cell lines for training/test can change (Table S1), moreover, they also have different number of doses, replicates, and the negative control used for normalization (Table S2). These 11 drugs are important enough to be studied in three different large-scale pharmacogenomics datasets and also they cover a wide range of drugs including chemotherapy agents, targeted therapeutics, FDA approved drugs, and experimental drugs (Table S1). It is important to note that the data we employed throughout this paper may be slightly different from the data accompanying the published studies because we obtained the data from the ORCESTRA platform, which hosts the integrated and standardized versions of these datasets, and because the datasets may have been updated by the study groups since their original publications.

### State-of-the-art in preclinical pharmacogenomics

We categorized the state-of-the-art predictors of drug sensitivity based on their input, output, and the pharmacogenomics datasets that they used for training and test (Figure 2). Gene expression was the most common input data type to predict drug sensitivity as it was determined to be the most effective data type in multiple studies [4,26,31,33]. However, some studies based on multi-omics data also demonstrated that adding other omics data types can improve the prediction performance [26,40]. For drug sensitivity, IC50 was the most common measure used. The cross-domain training approach was more common compared to the within-domain approach. Moreover, the majority of these methods were trained on GDSCv1 gene expression data. We also observed that incorporating drug structure, such as the SMILES representation of the drug molecule, is an emerging trend in the field. Since the goal of this study is investigating generalization in gene expression cell line-based predictors, we did not provide detailed descriptions of drug structure, interaction, adverse reaction, the type of clinical or PDX datasets that existing methods have employed and illustrated all of them under broad categories of “drugs”, “patients”, and “PDX”, respectively.

**Figure 2.**
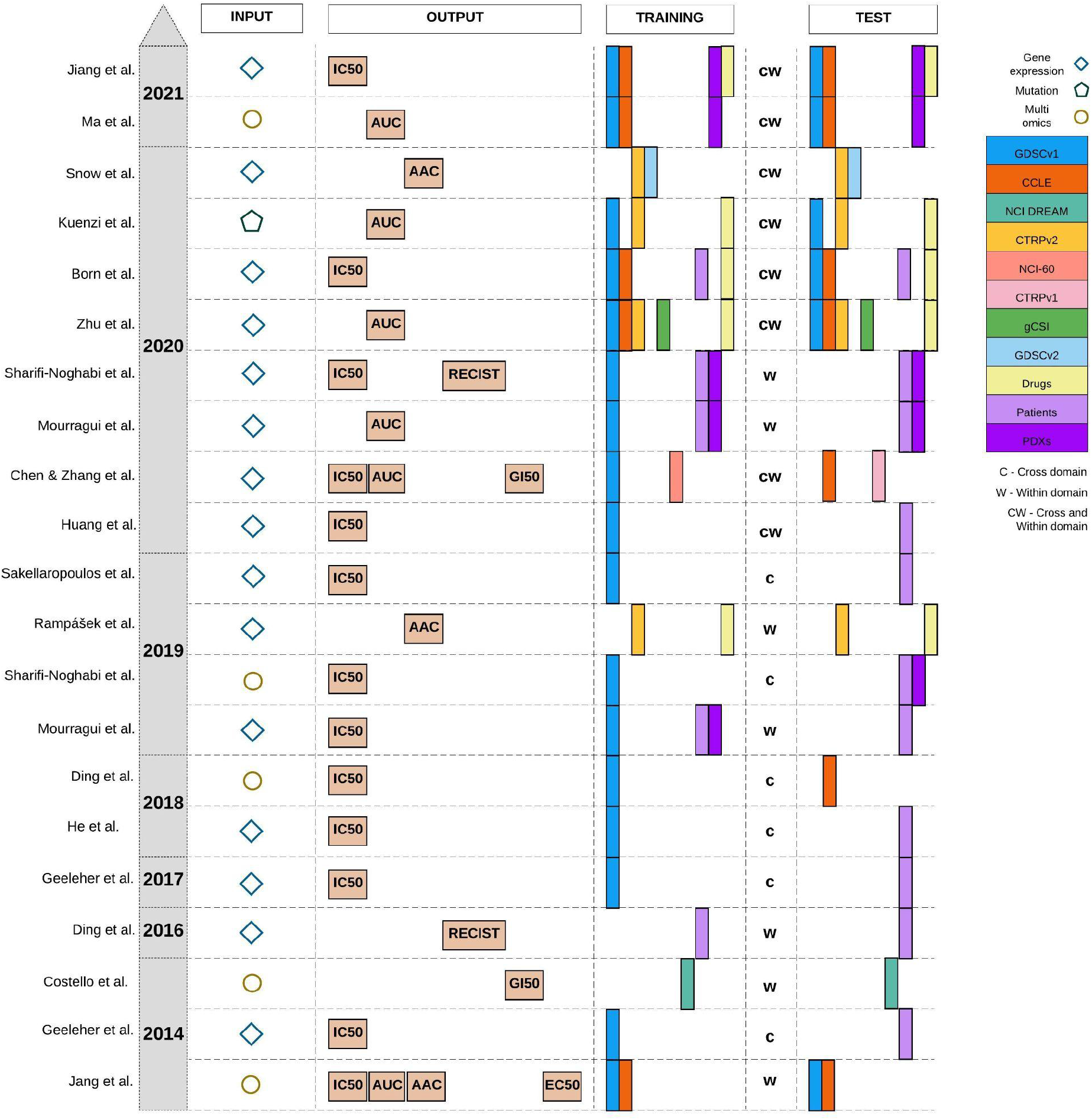
Published studies for drug sensitivity prediction. Gene expression is the most common molecular profile and IC50 is the most common pharmacological profile, but AAC/AUC has become more common in recent studies. GDSCv1 (originally named CGP) is the most common training dataset and the use of drug information for training has been more frequent in recent years. The cross-domain training approach denoted by “c” was more common compared to the within-domain approach denoted by “w”. When a method employs both of them, we denote it by “cw”.

### Univariable and Multivariable Analysis

To study the impact of model complexity on the performance and generalization of drug sensitivity predictors, we investigate a wide range of modeling approaches ranging from univariable modeling based on prospective biomarkers to highly complex and multivariable approaches such as Deep Neural Networks.

The univariable analysis consists of retrieving estimates for the association between previously studied prospective biomarkers and AAC or IC50 as measures of drug sensitivity. In this study, we focused on the prospective biomarkers of 11 drugs in common between the studied datasets as molecular features. A total of 35 unique prospective biomarkers were retrieved from literature for 8 drugs (out of 11) and used in the analysis (Table S3). The majority of these prospective biomarkers were based on gene expression but some of them were also based on mutation, copy number aberration, and gene fusion. Since the pharmacological datasets store genes as Ensembl gene ID, the biomarkers were mapped to the variant identifier using the Uniprot Retrieve/ID mapping tool (Table S4).

We explored four preprocessing approaches for IC50 including: using the raw IC50 values, log transformation of the values, truncating the values based on the concentration ranges of each study, and a combination of both. IC50 values can span several orders of magnitude, are bounded below by 0, and are often skewed. Log transformation tends to reduce the influence of outliers and brings the distribution of IC50 values closer to normal. Truncating based on predefined concentration ranges also reduces outliers, and reduces the influence of IC50 values which are extrapolated past the measured concentrations. These extrapolated values tend to be very sensitive to slight errors in estimation of the Hill Curve arising from noise in the measurements. AAC values were left unchanged.

Then, we used the PharmacoGx *drugSensitivitySig* function to compute estimates of the association between prospective biomarkers and drug sensitivity. For each measured gene this association is independently modelled using a linear regression model: 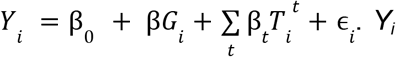. *Yi* denotes the measured drug sensitivity for sample *i*, *G_i_* denotes the measured gene expression for sample *i*, 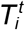 is an indicator variable for sample *i* belonging to tissue of origin *t*, ϵ_i_ is a random error term which is assumed to be normally distributed, and the βs are the estimated regression coefficients [49]. *Y* and *G* are scaled to have mean 0 and standard deviation of 1 prior to fitting the model, so that βs returned are standardized coefficients estimating the strength of the gene-drug association. The standardization facilitates comparison across genes and drugs, which may have very different scales and ranges of measured values. Note that this differs from a partial correlation in that scaling is done before adjusting for the covariates (tissues).

The multivariable analysis consists of making predictions of the drug sensitivity measures (AAC or IC50) given the level of expression of input gene features. Unlike univariable analysis which considers each biomarker at a time, multivariable analysis considers all of the input genes together. The goal of multivariable modeling is to learn a mapping function *Y* = *F_θ_ (X)* parametrized by one or more parameters θ that maps the input gene expression matrix *X^N×M^* to the drug sensitivity values *Y^N×1^*,where *N* is the number of samples and *M* is the number of input features. For all models considered in this paper, we employed the mean squared error as the loss function to optimize θ as follows: 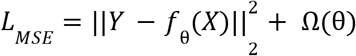, where Ω(.)denotes regularization. The regularization used was 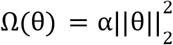 for Ridge Regression, 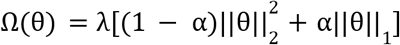 for Elastic Net, 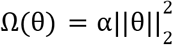 for the Deep Neural Networks, and no regularization was applied for the Random Forest models (λ and α are hyperparameters controlling the strength of the regularization). In addition to regularizing the norm of the parameters, the Deep Neural Networks were fit to the data with dropout and early stopping. More details on training and hyper-parameters of the final models corresponding to each method are provided in the supplementary material (Table S5).

### Within-Domain and Cross-Domain Analysis

To study the impact of data discrepancy on generalization of drug sensitivity predictors, we investigate two common approaches of within-domain and cross-domain. In within-domain analysis, the goal is to train and test models on the same dataset via cross-validation. The hypothesis is that if a model trained to predict sensitivity for a given drug cannot make accurate predictions for the same dataset (on the test splits), it is very unlikely that it generalizes to other datasets for the same drug. In cross-domain analysis, the goal is to train and test models on different datasets. The hypothesis is that models that demonstrate high performance in the within-domain should have better performance in cross-domain and models that perform poorly in within-domain should also perform poorly in cross-domain analysis. Intuitively, if there is enough predictive information in the training data for the given drug, the model should have a higher chance of generalizing to other datasets for the same drug (if the other datasets also have adequate predictive information).

### Experimental Design

We designed our experiments to justify the choice of training dataset (the input) and the measure of drug sensitivity (the output) as well as studying generalization of different models in within-domain and cross-domain analysis.

#### PGx Guideline experimental questions

IC50 is the most common measure of the drug sensitivity for machine learning (Figure 2) however it suffers from known limitations. By definition, the IC50 does not exist for any experiment where the maximum inhibition of growth is not at least 50%. Furthermore, IC50 estimation is unstable when there is not at least 1 point measured on each plateau of the curve [51], and the common technique to overcome this limitation, by setting IC50’s outside the measured range to the maximal tested concentration, effectively creates a right censored measurement and loses all differences in sensitivity between such experiments. Our hypothesis is that models trained to predict AAC should generalize better than those trained to predict IC50.

To answer the first PGx Guideline question on the best measure of drug sensitivity, we investigated a wide range of modeling approaches ranging from simple univariable models based on prospective biomarkers of each drug to more complex multivariable models based on Ridge Regression, Elastic Net, Random Forest, and Deep Neural Networks. We also compared CTRPv2 and GDSCv1 to determine which dataset is a better training dataset to build drug sensitivity predictors. We picked GDSCv1 as the competitor because it is the most common training dataset (Figure 2). GDSCv1 utilizes a different drug screening assay compared to the other datasets and for the majority of the drugs it has a smaller sample size [23]. Our hypothesis is that models trained on CTRPv2 are more generalizable because it utilizes the same assay as other datasets and also has a relatively larger sample size.

To answer the second question on generalization performance, we utilized the state-of-the-art cell line datasets (see Datasets section) in within-domain and cross-domain analysis. For within-domain analysis, we used 10-fold nested cross validation on CTRPv2, the largest dataset in our collection -- 9 folds for train and validation and the 10th fold for testing. For cross-domain analysis, we trained models on the CTRPv2 and tested them on the other cell line datasets (GDSCv2 and gCSI).

To answer the last question, we investigated the association between tissue type and model predictions when the model was trained with all available tissue types (solid and non-solid tissues) and when it was trained only on solid tissue types.

#### Evaluation

We employed different metrics in our analyses and experiments of the PGx Guidelines including Peason correlation, Spearman correlation, root mean squared error (RMSE), Jaccard index, and standardized regression coefficients. In all of the analyses, the Baseline performance indicates the correlation of cell lines in common between train and test dataset of that particular analysis. To summarize the figures, we also reported the average±standard deviation of the Pearson correlation over the 11 drugs in common (only in the main text).

#### Assessing stability of univariable feature rankings

To compare the rankings of univariable associations of gene expression (60662 genes) with drug response, we investigated the intersections between the top-K strongest associations (absolute value of standardized coefficient) for a range of K-values across GDSCv2, CTRPv2 and gCSI. We chose these datasets because they all shared the same drug response assay (CellTiter-Glo). We focus on the top-K rankings as weaker associations are more likely to be spurious due to the noise of the experiments, and therefore should not be expected to reproduce across datasets. For each drug, the Jaccard index between the three top-K lists was computed at each K. We then investigated two ways of measuring the stability of the top-K list across all three datasets. We first evaluated the minimum K (*K_min_*) for which the intersection was non-empty, which can be interpreted as measuring how many associations discovered in a single dataset would need to be tested across the other two datasets before a single hit is replicated. We also computed 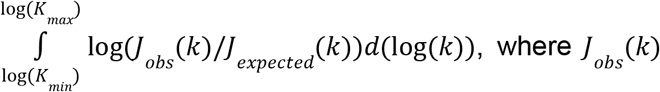, where *J_obs_(K)* was the Jaccard observed for the top-k intersection, and *J_expected_(k)*is the expected intersection if the rankings of the three lists were unrelated. The integral was estimated numerically on a grid of K values. For this integral, *K_min_* was chosen as the first value where *J_obs_(K)* was non-zero (consistent with above), removing values from the integrand that would otherwise be infinite (and negative). When comparing values for this integral, this can be seen as giving an unfair advantage to list-triplets which have a high *k_min_*, and therefore it should be evaluated in tandem with our first metric.

#### Implementation details

To ensure reproducibility of this study, we provide a detailed description of preprocessing, training, and evaluation. For all of the multivariable experiments, gene expression input data was normalized via z-score transformation using the parameters of the training dataset. Furthermore, to correct for the impact of tissue type, the one-hot encoded representation of it was added as an input feature to the normalized expression data after removing non-solid tissue types (except when the goal was to study the impact of non-solid samples) and those tissue types that were not available in the training data (CTRPv2).

We implemented the univariable analysis in R via the PharmacoGx package (version 2.0.5) [49]. The multivariable analyses were implemented in Python using the scikit-learn package (version 0.23.2). All of the hyper-parameter tunings were performed via grid search in nested 10-fold cross validation. We repeated the within-domain experiments 10 times and fixed the random seed for the cross-domain experiments and performed it once.

We implemented the deep neural networks in the Pytorch framework (version 1.4 cpu only) and used 10-fold cross validation and 100 trials of random search to select the best hyper-parameter settings for each drug.

In all of the analyses, we employed previously reported values of the hyper-parameters as our initial sets for each method and tuned to select the best setting for each method. More details on the considered values and the selected ones are provided in the supplementary material (Table S5).

#### Research reproducibility

All the data, code, and results employed and obtained in this study are publicly available for research reproducibility.

Code and supplementary tables/data: https://github.com/bhklab/PGx_Guidelines Data and models: https://zenodo.org/record/4642024#.YGCkbK9KiUl

## RESULTS

### Models trained to predict AAC outperforms those trained to predict IC50

Two common summary metrics have been used in the literature for summarizing dose-response experiments, the IC50 and the AAC. The IC50 is a measurement of potency, while the AAC can be interpreted as measuring an average of potency and maximal efficacy, or as a measure of the mean viability across the concentrations tested. While the IC50 is easily interpretable and is an absolute metric (unlike the AAC, which depends on the concentration range tested), the IC50 has some technical drawbacks which may make it difficult to use in training machine learning models. AAC/AUC is a normalized value between zero and one, but IC50 (the concentration) is not necessarily bounded and can be very small (close to zero) when samples are highly sensitive to a given drug or very large when they are highly resistant to a given drug. These issues make preprocessing of IC50 critical. Therefore, we investigated both methods to preprocess IC50, as one of the key measures of the drug sensitivity in previous studies, and then exploited univariable and multivariable analysis to compare these two metrics.

#### Univariable analysis using prospective biomarkers is not conclusive for preprocessing IC50

We employed univariable analysis of the prospective biomarkers on different preprocessing approaches for IC50 including, 1) estimating the associations using the raw IC50, estimating the associations using the truncated IC50, 3) estimating the association using log transformed IC50, and 4) estimating the association using log truncated IC50 values. We presented the standardized regression coefficients obtained from the univariable analysis for 8 drugs that we could obtain prospective biomarkers and highlight one biomarker for each drug (Figure 3A-Table S6). Across three datasets (CTRPv2, GDSCv2, and gCSI), we did not observe a clear winner for different ways of preprocessing IC50. This can be due to the fact that some of these biomarkers were based on gene expression data and some others based on mutation (Table S3) and this suggests further investigation.

**Figure 3.**
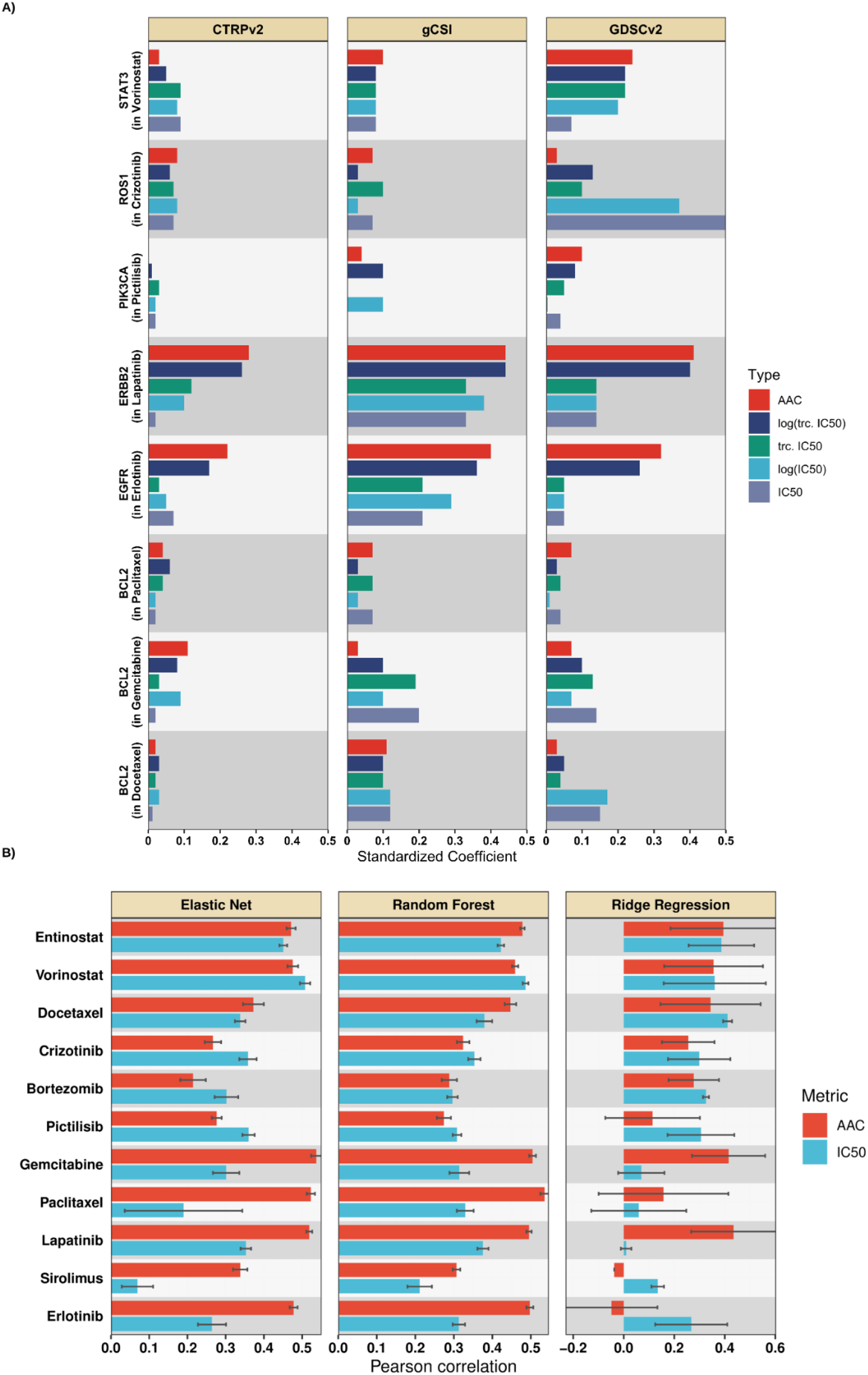
A) Comparison of AAC (Red) to different approaches of preprocessing IC50 via univariable analysis, tested on selected prospective biomarkers of the studied drugs in terms of the standardized coefficients. A single prospective biomarker per drug is shown (generally the marker with the strongest association), full results are available in supplementary data. B) Mean Pearson correlation (over 10 runs) for different multivariable methods in within-domain analysis trained to predict AAC or log-truncated log-truncated IC50. Multivariable within-domain analysis results indicated that AAC outperforms IC50 on average.

#### Univariable analysis using prospective biomarkers is not conclusive for AAC

We performed the same process of estimating associations for AAC and compared it to different ways of preprocessing IC50. Although for Lapatinib and Erlotinib AAC captures the associations between biomarkers and drug sensitivity more accurately compared to different approaches to preprocessing IC50, this pattern is not visible for other drugs (Figure 3A - Table S6). These results are not conclusive to compare AAC and IC50 (different ways to preprocess it) via the univariable analysis which also suggests further investigation.

#### Univariable cross-domain stability analysis suggests AAC and log-truncated IC50 produce most stable associations with drug response

For each gene with quantified expression (60662 genes) in the CTRPv2, gCSI and GDSCv2 datasets, we computed the strength of association with drug response for the 11 drugs in common across these datasets. We then ranked the associations by magnitude, and computed the Jaccard index for the 3-way intersection of the top K ranked univariable features for a range of K’s between 10 and 10,000 (with steps of 0.1 on a log10 scale). We examined two metrics: the first K at which the top-K intersection is non-empty (Figure S1A), and the integral of the observed Jaccard adjusted to the Jaccard expected by chance, on a log-log scale (Figure S1B). By both metrics, AAC produced the most stable or was tied for the most stable rankings of gene expression markers for the majority of the drugs (9/11 by both first non-empty K and integrated enrichment over null). Log-truncated IC50 similarly outperformed (or was ranked most stable or tied for most stable) the other transformations of IC50 (8/11 drugs by first non-empty K and 7/11 drugs by integrated enrichment over null). This suggests that AAC and log-truncated IC50 are better measures for multivariable analyses.

#### Multivariable within-domain analysis confirms that models trained to predict AAC outperforms those of log-truncated IC50

We compared Ridge Regression, Elastic Net, and Random Forest when trained on protein coding genes to predict AAC and log-truncated IC50, log IC50, truncated IC50, and raw IC50 in a within-domain analysis using CTRPv2.

The within-domain analysis using Ridge Regression and Elastic Net reconfirmed the stability analysis results that log-truncated IC50 outperforms the other approaches to preprocessing IC50 in terms of the studied metrics (Table S7). Interestingly, models for Docetaxel, Sirolimus, and Paclitaxel failed because of training to predict very large raw or log IC50 values but models for these drugs were successfully trained when using truncated IC50.

We observed that AAC achieved higher within-domain performance (Figure 3B -- Ridge Regression achieved 0.24±0.17 in AAC vs. 0.23±0.14 in IC50; Elastic Net achieved 0.4±0.17 in AAC vs. 0.31±0.12 in IC50; Random Forest achieved 0.41±0.1 in AAC vs. 0.33±0.07 in IC50). These results reconfirm the stability results that AAC is a better metric for drug sensitivity prediction compared to log-truncated IC50. We observed a similar pattern in the Spearman results (Figure S2) and RMSE (Table S8). In terms of Pearson and Spearman, seven drugs (out of 11) benefited from training to predict AAC instead of log-truncated IC50 in at least two different methods (out of three). These experimental results also align with the within-domain results in the previous study [48]. For simplicity, we refer to log-truncated IC50 as IC50 for the rest of the paper.

### Cross-domain analysis decreases generalization performance

To study the generalization capabilities of drug sensitivity predictors, we analyzed them in a cross-domain setting where the models are trained and tested on different cell line datasets. To perform this analysis, first we determined the most suitable training dataset.

#### Multivariable analysis reveals that models trained on CTRPv2 outperform those of GDSCv1 in generalization

As mentioned before, GDSCv1 is the most common training dataset for machine learning in pharmacogenomics. However, this dataset utilized the Syto60 assay in contrast to other major pharmacogenomics datasets that utilized the CellTiter Glo assay. We believe the difference in the drug screening assay influences the generalization capability of models because the Syto60 assay generates noisier drug response estimates [11]. To validate this, we trained two Ridge Regression models to predict AAC using protein coding genes: one trained on CTRPv2 and the other trained on GDSCv1 and then tested both of them on gCSI. We removed GDSCv2 from this analysis because of the overlap of molecular profiles with GDSCv1. We selected CTRPv2 because it is larger compared to gCSI which makes it naturally a more viable choice for training.

We observed that the model trained on CTRPv2 demonstrated significantly better performance in terms of the Pearson correlation compared to the model which was trained on GDSCv1 (Figure 4). To be more specific, on average (over 11 drugs), the model trained on CTRPv2 achieved the Pearson correlation of 0.4±0.21 (0.39±0.18 for IC50--Table S9), while the model trained on GDSCv1 achieved 0.26±0.16 (0.16±0.17 for IC50--Table S9). This suggests that agreement between the drug screening assay as well as sample size play significant roles in cross-domain generalization. We observed a similar pattern in the Spearman correlation (Figure S3) and RMSE (Table S9).

**Figure 4.**
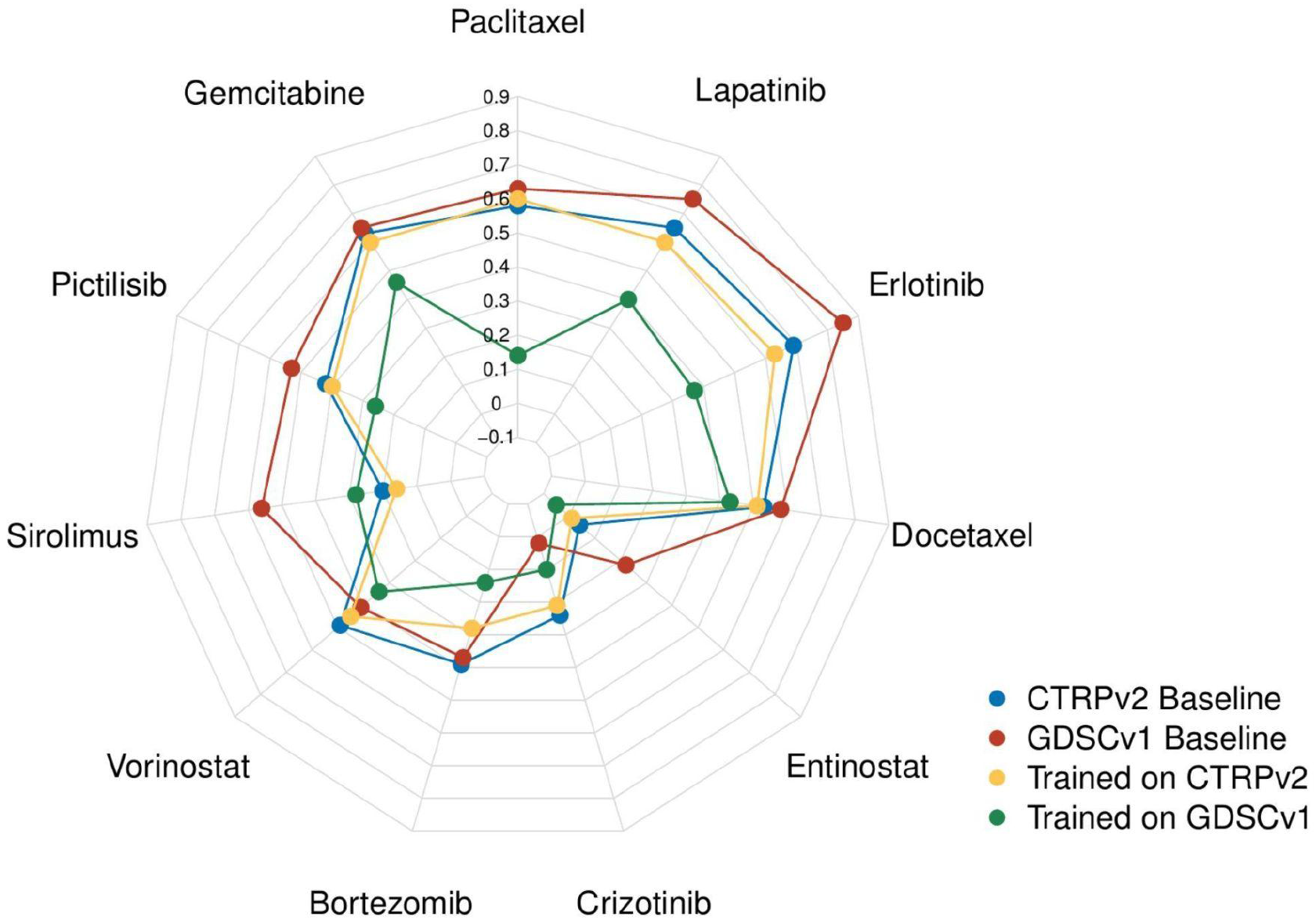
Comparison of CTRPv2 and GDSCv1 as training datasets for Ridge Regression models in terms of Pearson correlation in cross-domain analysis to predict AAC on gCSI. For the majority of the drugs, models trained on CTRPv2 (Yellow) achieved better performance compared to those trained on GDSCv1 (Green) even though GDSCv1 has higher baseline correlation (correlation of AAC values between cell lines in common between GDSCv1/CTRPv2 and gCSI -- CTRPv2 baseline in Blue and GDSCv1 baseline in Red).

#### Multivariable analysis reveals that the performance of models decreases when moving from within-domain to cross-domain analysis

To study the generalization of drug sensitivity predictors, we trained different models on CTRPv2 dataset and tested them of GDSCv2 and gCSI (Figure 5A). The cross-domain performance for the majority of the studied drugs is decreased significantly compared to the within-domain performance (Figure 5B-D). Elastic Net performance decreased from 0.4±0.17 (within-domain AAC) to 0.34±021 in GDSCv2 and 0.34±0.21 in gCSI (both in AAC- Figure 5C). Similarly, Random Forest decreased from 0.41±0.1 (within-domain AAC) to 0.33±0.2 in GDSCv2 and 0.35±0.21 in gCSI (both in AAC- Figure 5D). Ridge Regression demonstrated a different trend and increased from 0.24±0.17 (within-domain AAC) to 0.33±0.17 in GDSCv2 and 0.4±0.21 in gCSI (both in AAC- Figure 5A). This was due to some outlier predictions; Pearson correlation is sensitive to outliers and when we looked at the Spearman correlation results, the performance of Ridge Regression also decreased in cross-domain analysis (Figure S4 compared to Figure S2).

**Figure 5.**
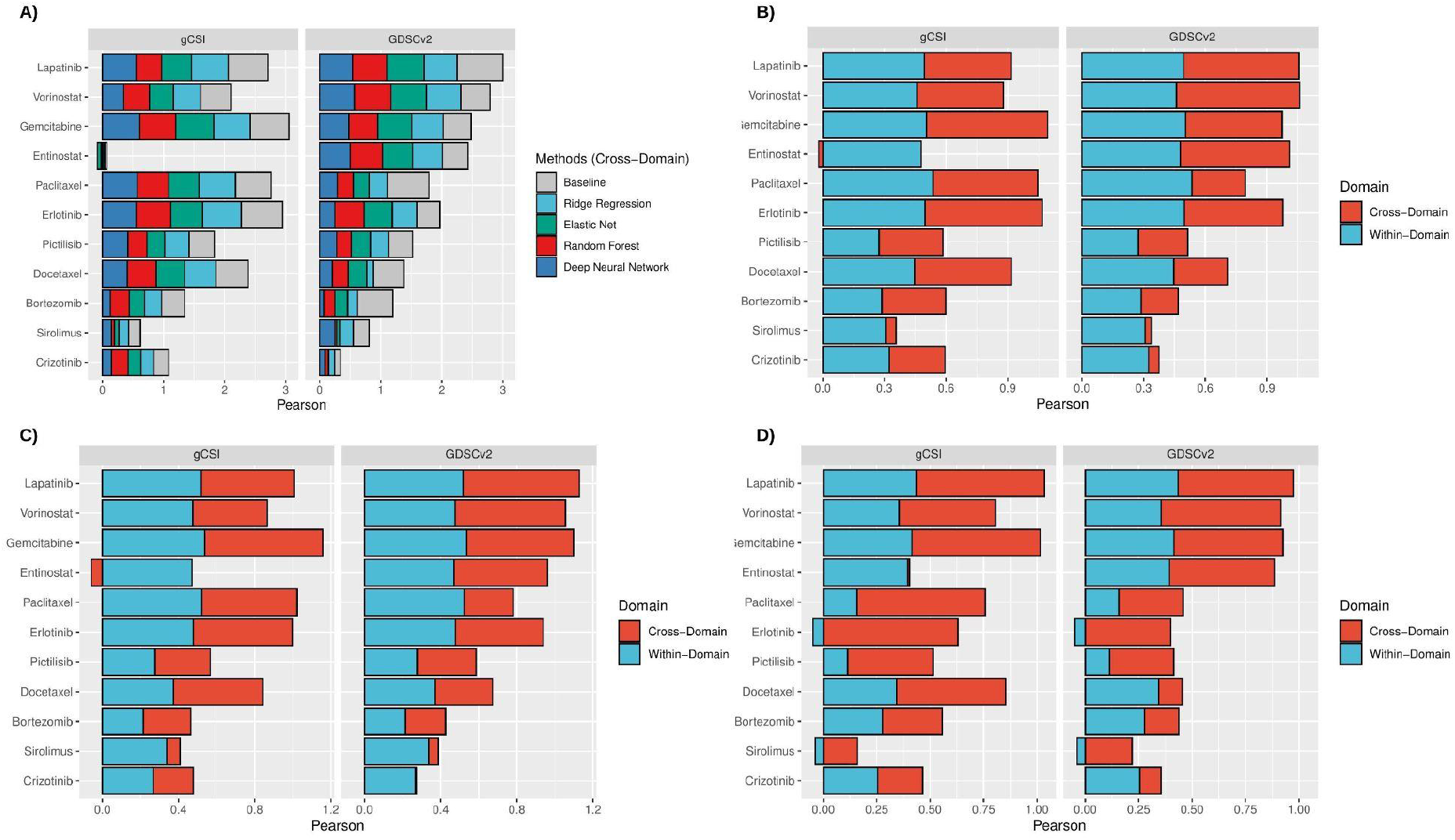
Comparison of multivariable methods in terms of Pearson correlation in cross-domain analysis trained on CTRPv2 to predict AAC, tested on GDSCv2 and gCSI (A). Comparison of the within-domain performance of Ridge Regression (B), Elastic Net (C), and Random Forest (D) to their cross-domain performance in terms of Pearson correlation.

These results suggest that even when the train and test data of a model utilized the same drug screening assay and were preprocessed similarly, it does not necessarily guarantee generalization. Moreover, the within-domain and cross-domain analysis together suggest that making models more complex improves the performance compared to the univariable analysis. For example, ERBB2 had an estimated association of 0.41 in GDSCv2 and 0.44 in gCSI, however, Elastic Net achieved 0.61 in GDSCv2 and Ridge Regression achieved 0.6 in gCSI which demonstrates the power of multivariable analysis. Finally, the cross-domain analysis using multivariable methods also reconfirmed that AAC is a better metric compared to IC50 (Table S10) because datasets are more consistent on AAC (baseline correlations of 0.45±0.18 in GDSCv2 and 0.43±0.21 in gCSI for AAC in contrast to 0.38±0.27 and 0.42±0.13 in IC50, respectively) and methods are more accurate (for example, in gCSI, Ridge Regression achieved 0.4±0.21 in AAC vs. 0.36±0.18 in IC50; Elastic Net achieved 0.34±0.21 in AAC vs. 0.33±0.18 in IC50; DNN achieved 0.35±0.2 in AAC vs. 0.32±0.21 in IC50; Random Forest achieved 0.35±0.2 in AAC vs. 0.37±0.16 in IC50). We observed a similar pattern in the Spearman correlation (Figure S3A-D). We also observed similar patterns in both Pearson and Spearman results when comparing the best performing within-domain model to the best performing cross-domain model for each drug (Figure S4E-F).

While employing gCSI offers more robust comparison across multiple datasets, it also limits the number of drugs that we can study because this dataset is only screened with 16 drugs. To study the cross-domain performance on more drugs, we limited our focus to 70 drugs in common between CTRPv2 and GDSCv2. We trained three models using Ridge Regression, Elastic Net, and Random Forest for each drug using CTRPv2 to predict AAC and IC50 separately and tested the performance in terms of Pearson, Spearman, and RMSE on GDSCv2. We removed those drugs that have less than 100 samples for training or 50 samples for tests or had a failed model for AAC or IC50 to ensure a fair comparison.

We observed that the results are fairly competitive between models trained to predict AAC and those of IC50 when we only focus on one dataset. However, RMSE shows that AAC outperforms IC50 which can be due to larger magnitude of IC50 values as opposed to AAC (Table S11).

#### Non-solid tissue types influence the performance of models

The majority of pharmacogenomics datasets are pan-cancer with solid and non-solid tissue types. We studied the molecular and pharmacological profiles of non-solid tissues (hematopoietic and lymphoid tissue types) in CTRPv2, GDSCv2, gCSI. We observed that the sensitivity outcome (AAC) in non-solid samples is significantly different compared to solid samples and they tend to be more sensitive than solid samples (Figure S5A-C) which aligns with the previous studies [1,52,53]. Similarly, the non-solid samples also clustered differently compared to solid samples (Figure S5D-F). These results raise the question of whether including both liquid and solid lines in the training set is beneficial for learning models to predict drug sensitivity.

To answer this, we trained two Ridge Regression models to predict AAC using protein coding genes as follows: one trained on all samples (solid and non-solid together) in CTRPv2, and the other one trained only on solid samples (non-solid samples were removed) in CTRPv2. We measured the associations between the predictions and the binary status of tissue type (solid vs. non-solid) in GDSCv2 and gCSI using the area under precision-recall curve (AUPR). The predictions of the model which was trained on all samples demonstrated a very high AUPR compared to the model that was only trained on solid samples. This suggests that by including non-solid samples, models predict the tissue type rather than the drug sensitivity itself (Figure 6). To confirm this, we trained another Ridge Regression model after removing a random subset of solid samples with the same size as the non-solid samples to make sure that the observed result was not because of sample size (we repeated the random selection 10 times and reported the average value). We also reported the Baseline AUPR which indicates the ground truth association between actual AAC and the binary status of tissue type. We observed that models that were trained on all samples (including solid and non-solid tissues) and the one with a random subset removed, had the highest AUPR in both GDSCv2 and gCSI for the majority of the drugs compared to the model that was trained only on solid samples (Figure 6). This confirms previous results that the difference in molecular profiles and drug sensitivity of non-solid samples have a negative impact on the drug sensitivity prediction task and it is crucial to remove all non-solid tissue types before any machine learning modeling.

**Figure 6.**
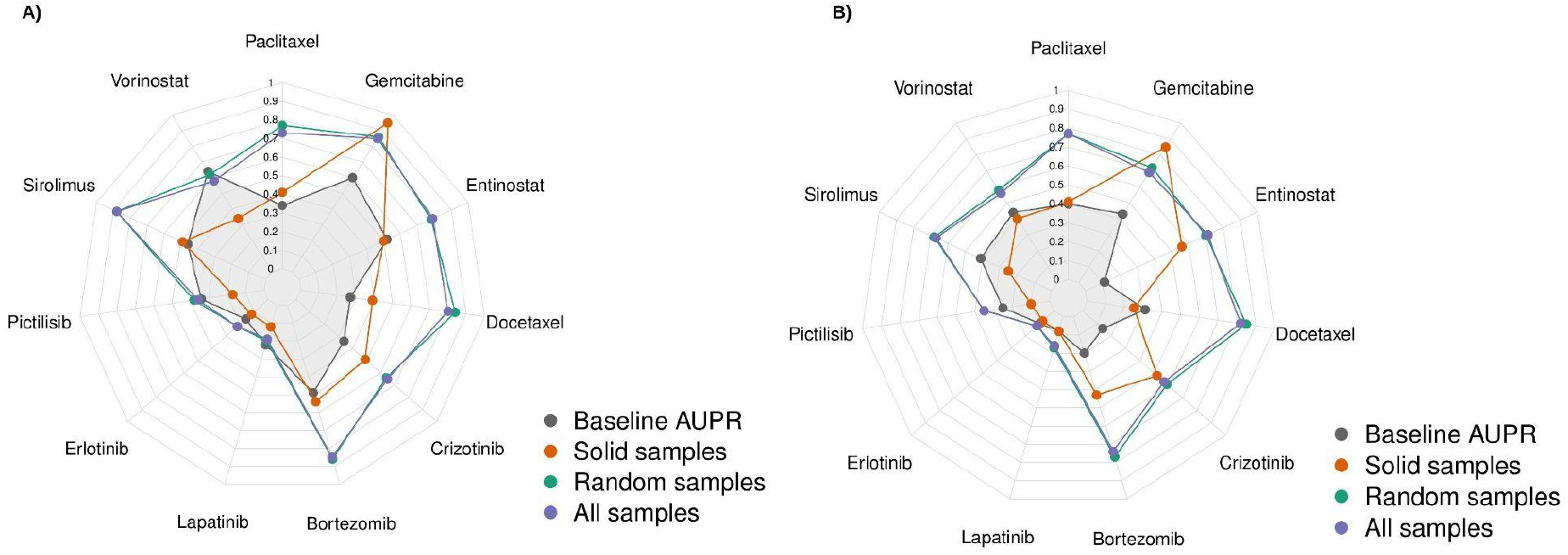
The association between predictions of a Ridge Regression model trained on CTRPv2 to predict AAC and tissue binary type status of GDSCv2 (A) and gCSI (B) in terms of AUPR. Training on cell lines with solid (Red) tissue types had lowest AUPR compared to training on all samples (solid and non-solid--Purple) and training on a randomly selected subset with the same size as solid samples (average AUPR over 10 different subsets--Green). The results confirmed that removing non-solid samples decreases the association of predictions with tissue type.

We also studied the impact of different tissue types on the performance by comparing three scenarios: 1) training a Ridge Regression model on solid and non-solid tissues combined and testing it on solid tissues only, non-solid tissues only, and all tissues combined. 2) training a Ridge Regression model on solid tissues only and testing it on solid tissues only, non-solid tissues only, and all tissues combined. Finally, 3) training a Ridge Regression model on non-solid tissues only and testing it on solid tissues only, non-solid tissues only, and all tissues combined. For each analysis we used CTRPv2 as the training dataset to predict AAC, and tested the model on GDSCv2 and gCSI and reported the results in terms of Pearson, Spearman, and RMSE. For each scenario, the dataset with the largest sample size was downsampled to have the same size as the smaller one 10 times to control for this factor. We observed that on average (over 10 runs), models trained on non-solid tissues had the best performance when tested on non-solid tissue types and similarly, models trained on solid tissues had the best performance when tested on solid tissues. This reconfirms the importance of removing non-solid tissue types from the training data for solid tissues (Table S12).

## DISCUSSION

In this study, we investigated the fundamental challenges of developing machine learning models to predict drug sensitivity from cell line pharmacogenomics data. We named our study PGx Guidelines because we believe that the answers to these questions provide actionable guidelines for developing predictors of drug sensitivity.

The guidelines show that the performance of machine learning models decreases when moving from within-domain multivariable modeling to cross-domain multivariable modeling. This is particularly important because it shows that models face generalization difficulties when trained and tested on cell line datasets with comparable molecular and pharmacological profiles. Consequently, such models are highly unlikely to generalize to clinical samples when they fail to generalize to (more similar) preclinical samples.

The PGx guidelines also demonstrate that the AAC is a more suitable measure of drug sensitivity as opposed to the IC50, and CTRPv2 is a more suitable training dataset as opposed to GDSCv1 due to the larger sample size available with RNA sequencing and the difference in the drug screening assay. This is particularly important because employing IC50 and GDSCv1 is currently the trend in machine learning for drug sensitivity prediction.

Finally, The PGx guidelines demonstrate the necessity of removing non-solid tissue types from the datasets before any modeling which is often not considered in training models. This is especially true for post-hoc interpretation of feature importance. Our results suggest that models trained on a mixture of these two tissue types primarily learn to predict non-solid tumour status, bringing into question whether importance scores will be relevant to the task of drug sensitivity prediction.

Our goal was not to provide a comparative study -- train models to achieve the highest possible performance -- in terms of method; we selected some of the most basic available methods to focus on the importance of data for drug sensitivity prediction. The reported results can likely be improved by investing more time on hyper-parameter tuning or adopting more complex training schemes or objective functions (particularly for Deep Neural Networks). However, our experiments shed light on some of the current issues with machine learning for drug sensitivity prediction.

Although Ridge Regression and Random Forest showed slightly better performance compared to Elastic Net, overall, these methods showed a competitive performance. We focused on Ridge Regression for the majority of the analyses of PGx Guidelines because it is less sensitive to the setting of hyper-parameters than other methods, in particular Deep Neural Networks, which have the highest number of hyper-parameters and are more sensitive to the choice of their values. Therefore, we included Deep Neural Networks for cross-domain analysis but did not consider them for other experiments. We also did not perform within-domain analysis for Deep Neural Networks because of the limited sample size of the nested cross validation for the high number of parameters of each network and the early stopping regularization which makes the comparison difficult for within-domain analysis. We also utilized feature selection to reduce the input dimensionality (number of genes) and tried focusing only on the L1000 landmark genes [54], or focusing on top genes selected by the Minimum Redundancy--Maximum Relevance (mRMR) method [55]. However, we did not observe any significant difference (Figure S6 and Table S13).

Some of the major limitations of this study are as follows: we assumed a similar concentration range across the studied datasets, this can be an important factor in generalization but the problem is that by focusing on the samples with the same range, we will not have enough samples to train models especially given the high dimensionality of the data. We believe this is a very important factor that should be considered when more samples are available [56,57]. Similarly, we did not consider multi-omics data because comparable omics data types (mutation, proteomics, copy number aberration, etc.) are not available in the studied pharmacogenomics datasets to investigate the impact of multi-omics data on cross-domain generalization [33,40]. For PGx Guidelines, we only focused on monotherapy models and did not investigate multi-output or multi-task learning, or drug combination which can be promising future direction, similar to incorporating the chemical structure representation of the drugs as input. Also, we did not consider the pathway transformation of the genes as input features and employed genes themselves. Such transformations can improve the prediction performance [33]. We studied gene expression-based predictors, and our approach can be replicated for other omics data types when more data is available, and for other gene/feature representations such as pathway representation which is covered in other studies [33]

Investigating the source of the performance drop occurring when moving to cross-domain analysis is outside the scope of this current study. However, there have been extensive studies into inconsistencies between drug screening experiments on the same cell lines, and the reasons why they arise [11,16,21,58]. These studies have shown that some inconsistency can be explained by differences in experimental protocols, including: choices of drug concentration range and number of tested points, cell seeding densities, timepoints for measuring viability, cell viability assays used, number of technical and biological replicates, growth media and different choices for positive and negative controls (we summarize a subset of these variables for the studies used in Supplementary Table S2) [11,17,58]. Genetic drift in cell lines, and different cell line doubling times between labs has also been shown to affect drug sensitivity measures such as the IC50 and AAC [58–61]. Finally, technical sources of variation, even with identical (or as close as possible) protocols, both in executing the experiments and subsequent analysis have been shown to lead to considerable variability between labs [21]. Importantly, Niepel et al. found that inconsistencies between labs often arise when experimental differences interact with biologically meaningful variation, meaning that particular cell lines or drugs may be strongly affected by differing experimental decisions while others are not [21]. It is also important to remember that IC50 and AAC are complex phenotypes which can only be measured indirectly through accessing a dose-response curve and fitting a model to this data. While in our study we have removed variation arising from choices of different curve estimation methods, all the sources of variability discussed above affect each measured point on these curves and the error in the measurement. This means that the model used to calculate IC50 and AAC will unavoidably have different bias and variance characteristics between studies. Overall, measuring and analyzing drug response data in cell lines is technically complex, and given that there is no consensus experimental and analytical protocol e, our findings reinforce the importance of checking performance across cell line domains to truly understand the robustness and generalizability of machine learning models in this field.

In summary, the key takeaways of PGx Guidelines are:

- Models tend to be more accurate when trained to predict AAC rather than trained to predict IC50.
- If IC50 is used for the modeling, truncating the IC50 values after logarithmic transformation yields more predictive models.
- Models trained on CTRPv2 to predict AAC tend to be more accurate than those trained on GDSCv1 which is partly due to the size of the dataset and the use of different cell viability assay (which considerably reduce the consistency across datasets)
- In pan-cancer datasets, our results indicate that it is advisable to stratify the analysis by tissue types, in particular solid vs non-solid cancer cell lines. It is important to note that we did not perform a comprehensive comparison of all existing methods and some modeling strategies may be able to leverage the difference between solid and non-solid cancer cell lines to develop more generalizable models.
- To evaluate the predictive performance, only looking at one metric might not be sufficient and it is more reliable to study multiple metrics.
- We suggest a modeling path that starts with simple analysis using one gene (biomarker), then performs multivariable modeling within one dataset, and eventually performs multivariable modeling across multiple datasets. We note that testing on one cell line dataset only does not even give an adequate measure of model performance on another cell line dataset.

It is important to note that our guidelines do not cover best practices to choose specific methods or types of input data. We refer interested readers to previously published literature that has extensively explored these topics [26,33,48,56,61].

## ACKNOWLEDGEMENTS

We would like to thank Hossein Asghari, Colin C. Collins (Vancouver Prostate Centre), and Ian Smith (Princess Margaret Cancer Centre) for their kind support.

---

**Figure S1.**
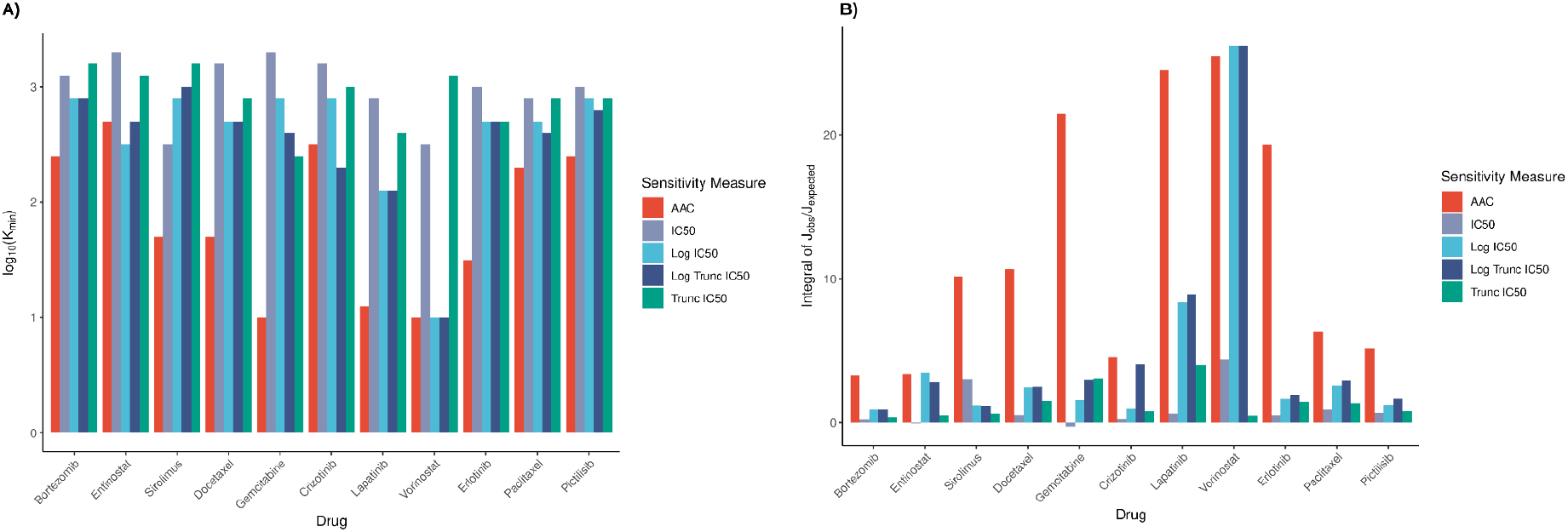
Assessment of consistency for the top-K univariable associations across CTPRv2, GDSCv2 and gCSI, using different measures for drug sensitivity. The log10 of minimum value of K for which there is a non-empty intersection (K_min_) are plotted for each drug and each sensitivity measure (A), with lower numbers signifying greater consistency. AAC and Log Truncated IC50 are most often the most consistent measures using this metric. The integral of the observed Jaccard divided by the expected Jaccard on a log-log scale (B) reveals the same results, with AAC and Log Truncated IC50 again most often leading to the most consistent univariable associations.

**Figure S2.**
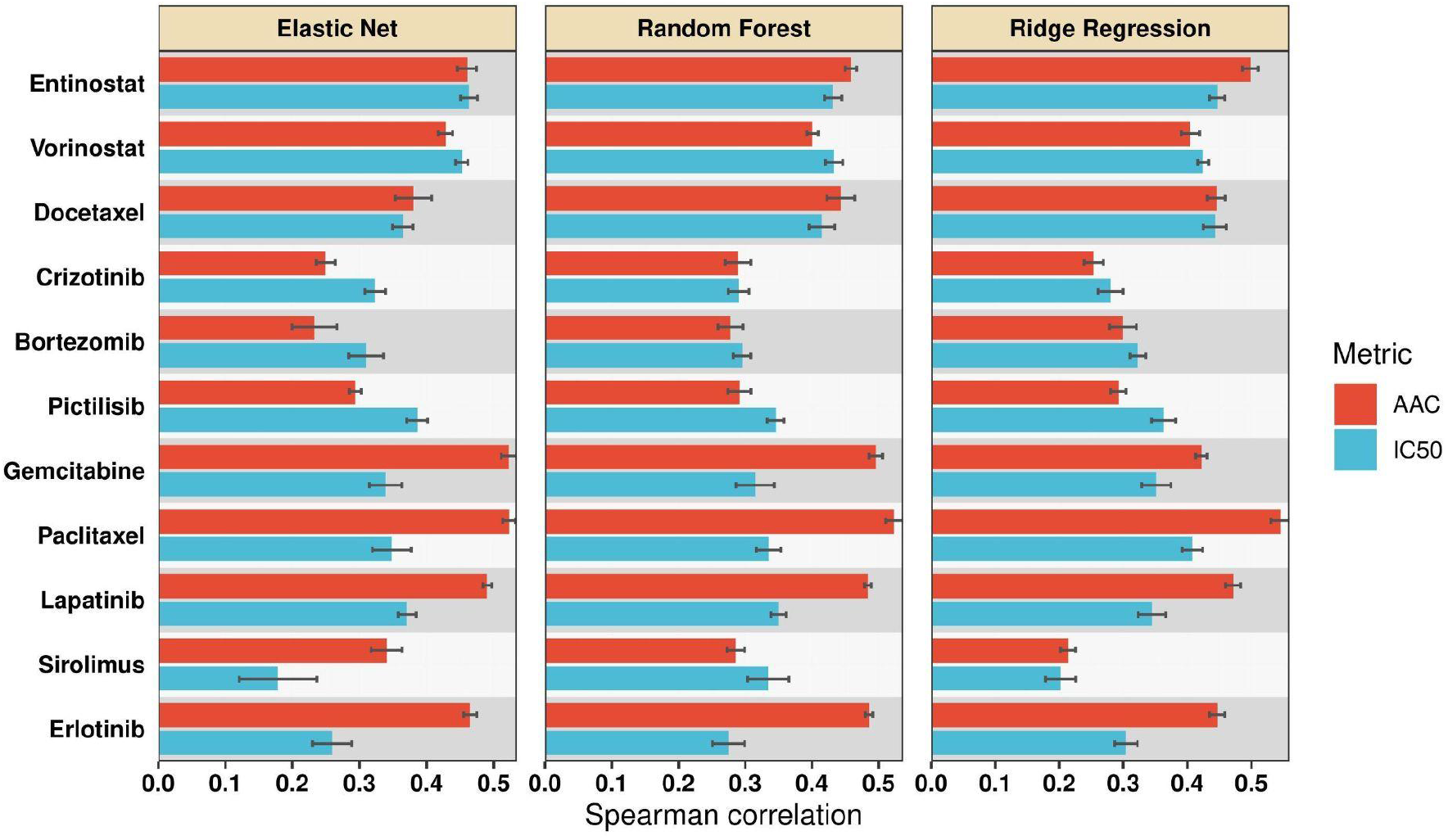
Mean Spearman correlation (over 10 runs) for different multivariable methods in within-domain analysis trained to predict AAC or IC50. The results indicated that AAC outperforms IC50.

**Figure S3.**
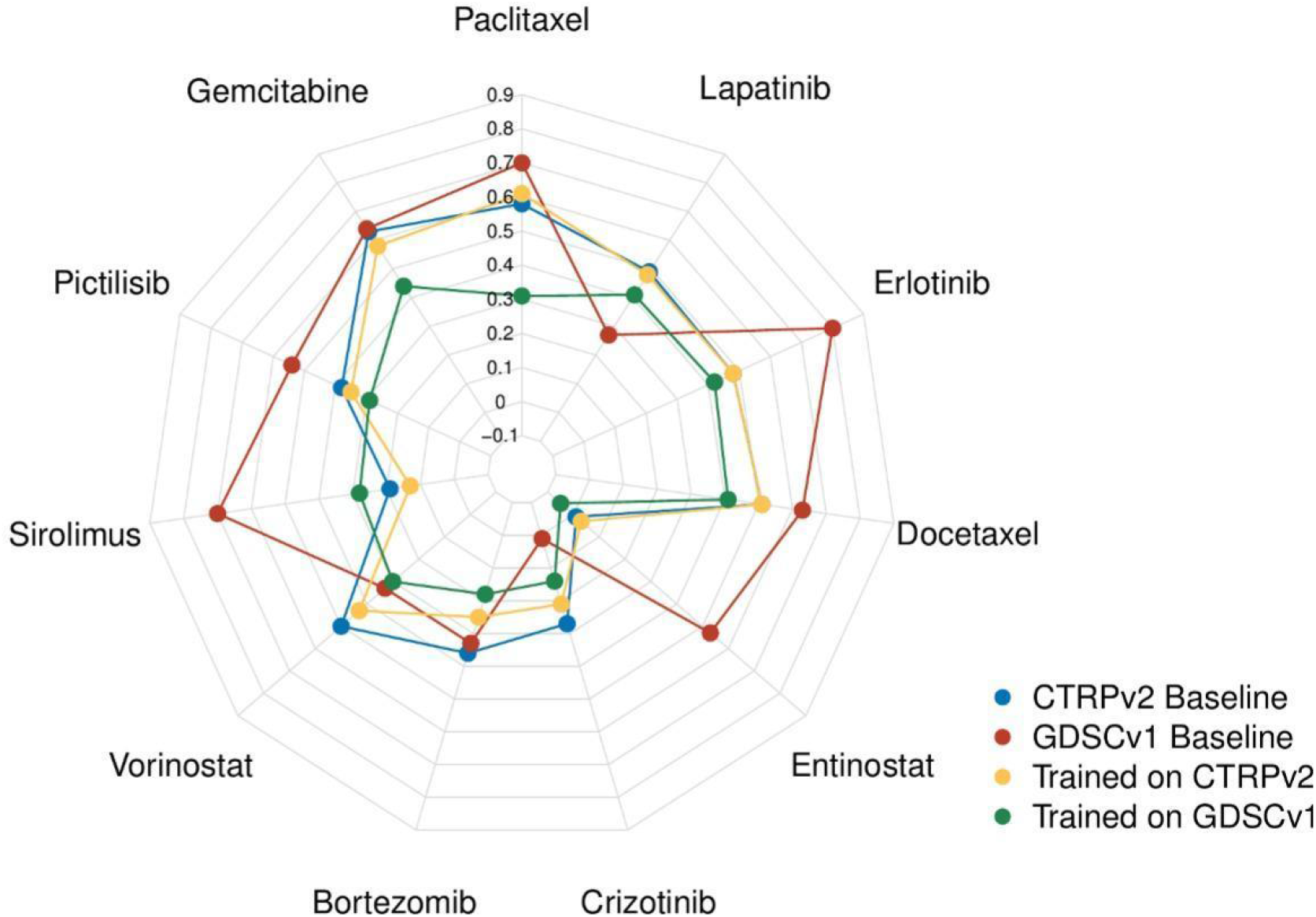
Comparison of CTRPv2 and GDSCv1 as training datasets for Ridge Regression models in terms of Spearman correlation in cross-domain analysis to predict AAC on gCSI. For the majority of the drugs, models trained on CTRPv2 (Yellow) achieved better performance compared to those trained on GDSCv1 (Green) even though GDSCv1 has higher baseline correlation (correlation of AAC values between cell lines in common between GDSCv1/CTRPv2 and gCSI--CTRPv2 baseline in Blue and GDSCv1 baseline in Red).

**Figure S4.**
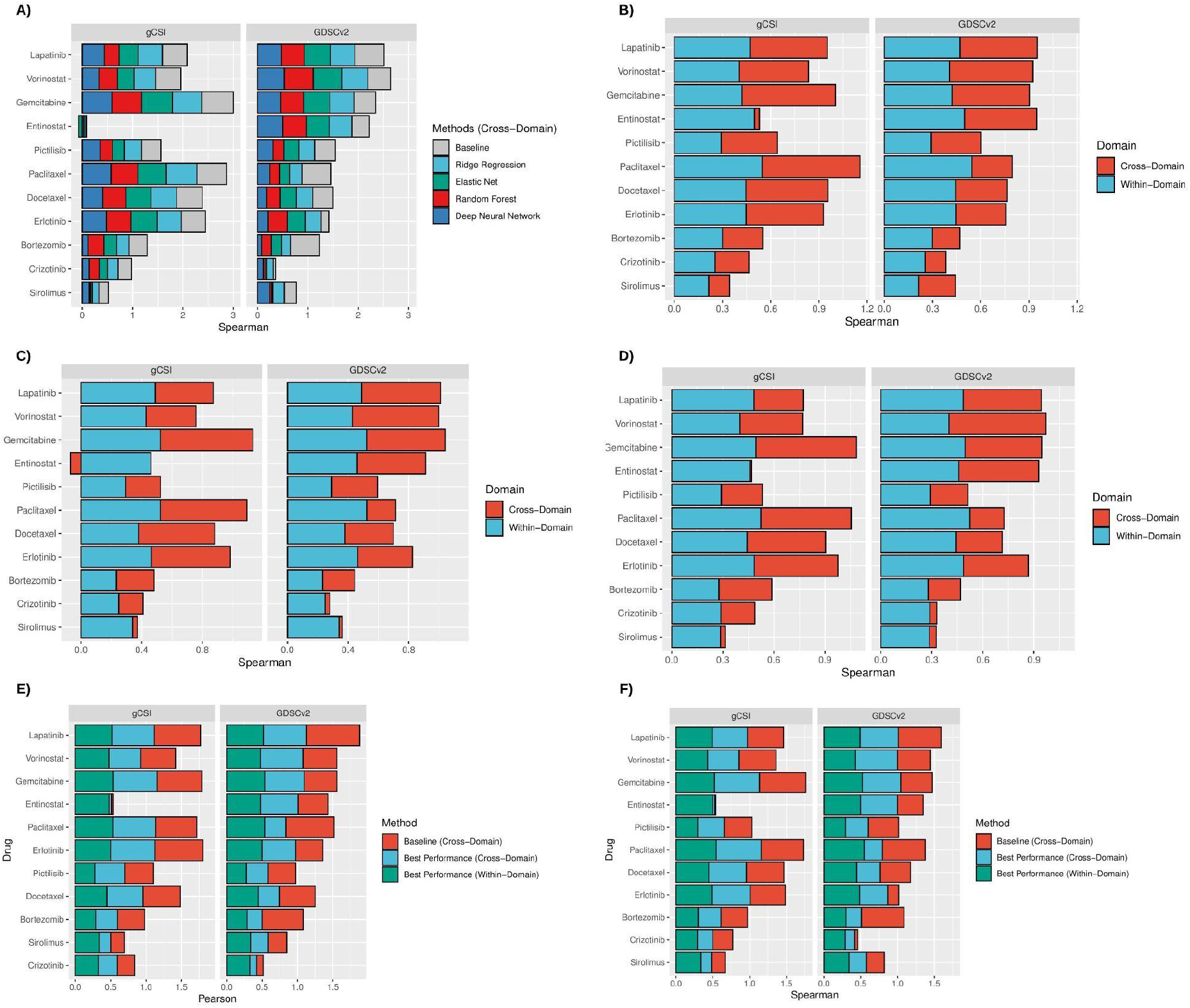
Comparison of multivariable methods in terms of Spearman correlation in cross-domain analysis for AAC.

**Figure S5.**
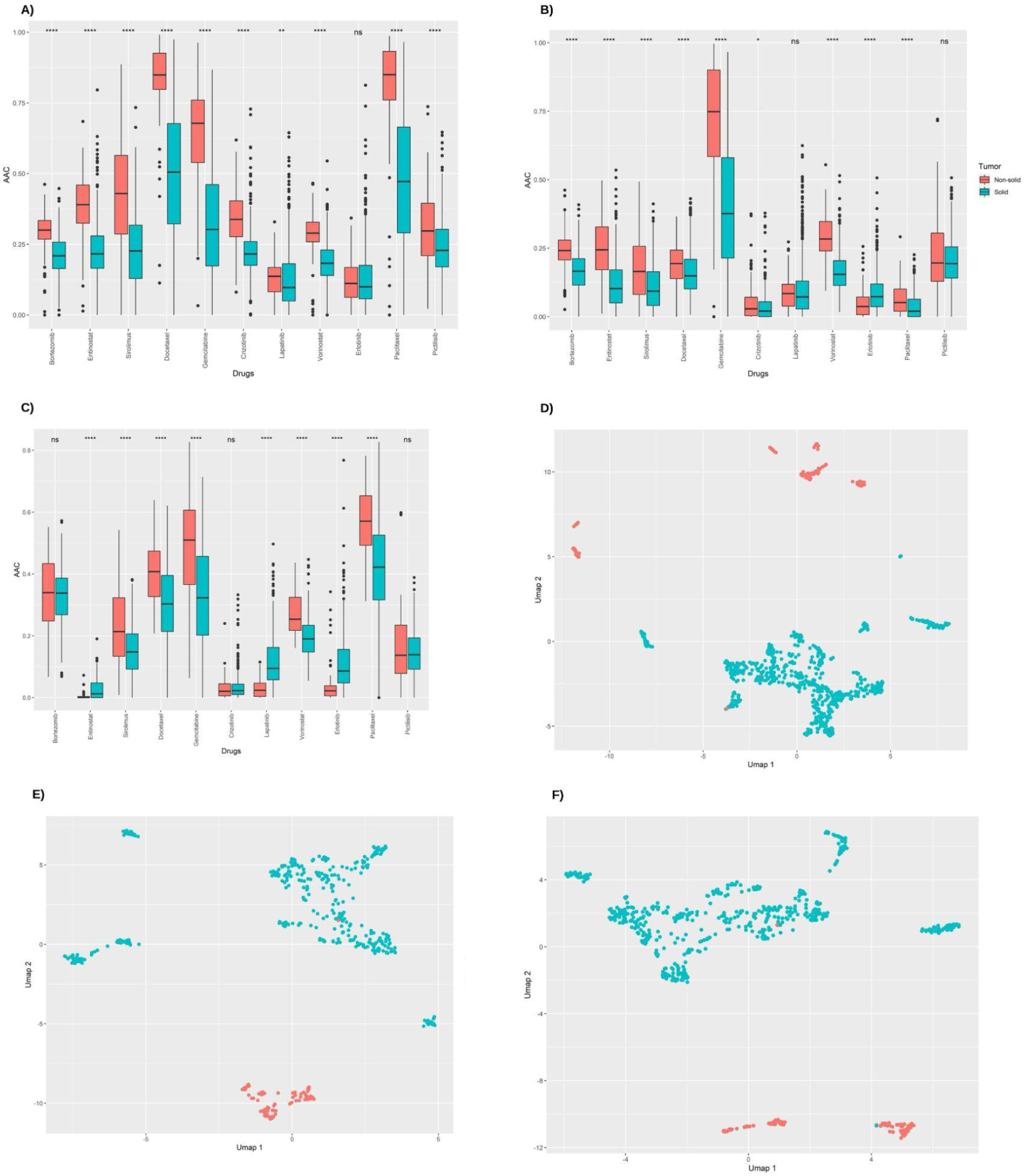
Comparison of the AAC values of solid (Blue) and non-solid (Red) samples across CTRPv2 (A), GDSCv2 (B), and gCSI (C) for 11 drugs in common with them. Comparison of the molecular profiles of solid (Blue) and non-solid (Red) samples via UMAP plot across CTRPv2 (D), GDSCv2 (E), and gCSI (F).

## Feature Selection

To see if reducing the number of input genes improves the prediction performance, we investigated two feature selection approaches: II) focusing on L1000 landmark genes [54], and focusing on top 200 genes selected by the Minimum Redundancy--Maximum Relevance (mRMR) method [55]. The idea of mRMR method is to select a subset of features (genes) that have high correlations with the output (drug sensitivity) and minimum correlations among themselves. We trained Ridge Regression models on CTRPv2 using the reduced features and tested them on GDSCv2 and gCSI in a cross-domain fashion. The experiments of L1000 landmark genes (961 genes were common across applied datasets) and mRMR top 200 genes were implemented in R via the mRMRe (version 2.1.0) and Caret packages (version 6.0.86). On average, we did not observe a significant difference between feature selection cross-domain results and the cross-domain results obtained from protein coding genes for AAC (Figure S6).

---

**Figure S6.**
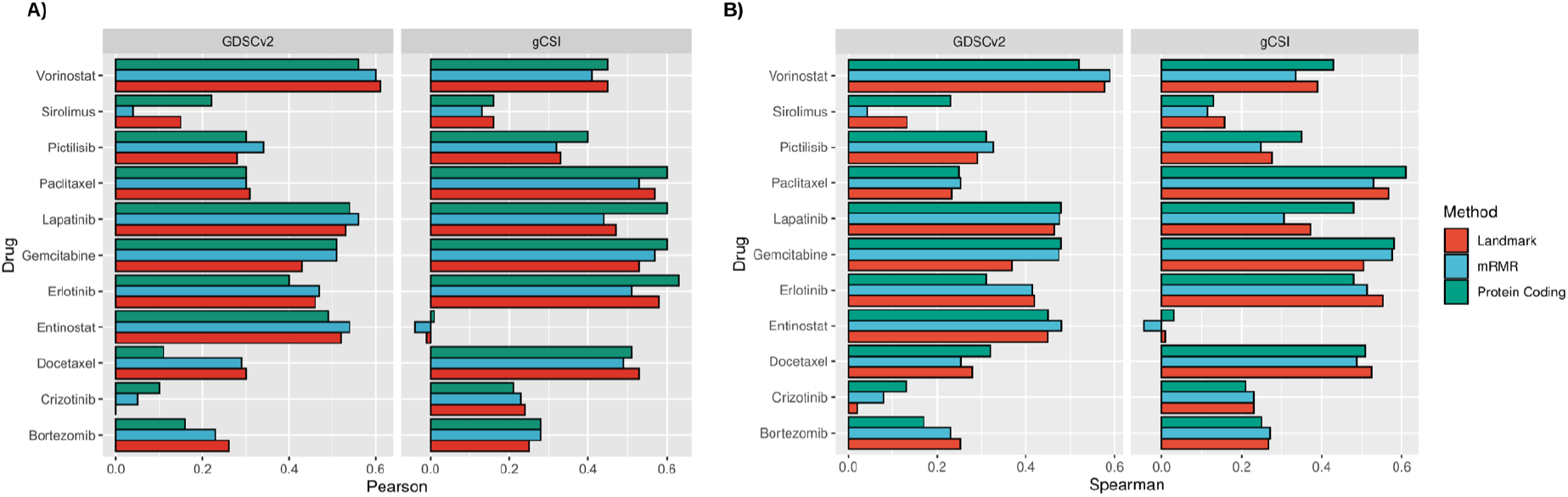
Comparison of top 200 genes selected by mRMR (Blue) and L1000 landmark genes (Red) with protein coding genes (Green) on GDSCv2 and gCSI for AAC in terms of Pearson correlation (A) and Spearman correlation (B).

